# Co-administration of intranasal parainfluenza virus vaccines expressing antigenically distinct SARS-CoV-2 S antigens elicits broad and durable immunity in hamsters

**DOI:** 10.1101/2025.10.22.683849

**Authors:** Hong-Su Park, Yumiko Matsuoka, Celia Santos, Eleanor F. Duncan, Jaclyn A. Kaiser, Lijuan Yang, Cindy Luongo, Xueqiao Liu, I-Ting Teng, Peter D. Kwong, Ursula J. Buchholz, Cyril Le Nouën

## Abstract

Intranasal COVID-19 vaccines with the ability to induce broad and durable mucosal and systemic immunity would be useful as stand-alone vaccines or in combination with injectable vaccines. Here, we evaluated in the hamster model the immunogenicity, breadth of immunity, and durability of protection elicited by co-administration of two live-attenuated bovine/human parainfluenza virus type 3 (B/HPIV3) vectors expressing antigenically distinct prefusion stabilized S proteins of the ancestral SARS-CoV-2 isolate (B/HPIV3/S-6P) or the Omicron/BA.5 variant (B/HPIV3/S-BA.5-2P). These vectors are being developed as bivalent pediatric vaccines against HPIV3 and SARS-CoV-2 and are based on bovine PIV3 with the fusion (F) and hemagglutinin-neuraminidase (HN) glycoproteins replaced by those of human PIV3. To broaden the S-specific antibody response, we evaluated co-administration of these B/HPIV3 S-expressing vectors. Each B/HPIV3 S-expressing vector induced robust serum anti-S IgG and IgA antibody levels to the antigen-matched S protein that were sustained for at least five months. Co-administration increased the breadth of the S-specific antibody response, spanning the antigenic breadths of the response elicited by each B/HPIV3 S-expressing vector individually. Animals that had received the mixture of vectors developed neutralizing antibodies to ancestral as well as recently circulating SARS-CoV-2 strains. Hamsters immunized intranasally were protected against Omicron/BA.5 challenge 5 months after immunization, with no weight loss, SARS-CoV-2 challenge virus replication, or increase in host inflammatory cytokines in the upper and lower airways detectable after the challenge, indicating durable protection. Thus, intranasal co-administration of live-attenuated B/HPIV3 expressing antigenically distinct S proteins induced broad and durable antibody responses and long-term protection against Omicron/BA.5 challenge. This approach warrants further development and may better protect against emerging SARS-CoV-2 variants.

**Author summary:** Current SARS-CoV-2 vaccines protect against severe disease but are less efficient at blocking infection. Intranasal SARS-CoV-2 vaccines, however, have been shown to induce local immunity that better blocks infection at the nasal portal of entry and have been proposed as booster vaccines to induce better protection against SARS-CoV-2 variants. In the present study, we evaluated in hamsters an intranasal live-attenuated chimeric bovine/human parainfluenza virus type 3 (B/HPIV3) as a bivalent pediatric vaccine against PIV3 and SARS-CoV-2. Co-administration in a single intranasal dose of two B/HPIV3 vectors, one expressing the spike protein S from the antigenically distinct ancestral Wuhan-Hu-1 strain and one expressing S of Omicron/BA.5, induced broad and durable serum anti-S IgG and IgA antibody responses that remained strong for at least five months. Hamsters challenged with the Omicron/BA.5 strain five months after immunization were protected from weight loss, inflammatory responses, and challenge virus replication in the upper and lower airways after challenge. Thus, in the hamster model, intranasal immunization with live-attenuated B/HPIV3 expressing SARS-CoV-2 S can provide durable protection for several months, and combining two antigenically distinct B/HPIV3 S-expressing vectors substantially broadens the antibody response. This mucosal immunization approach may better protect against infection from emerging SARS-CoV-2 variants.

## Introduction

Injectable SARS-CoV-2 vaccines including mRNA-based vaccines have been highly effective in reducing COVID-19 hospitalization and mortality. However, they remain limited in their ability to prevent SARS-CoV-2 infection and replication in the upper respiratory tract, and their efficacy in blocking viral transmission and spread is low. Immunity induced by mRNA vaccines or SARS-CoV-2 infection wanes rapidly over 4 to 6 months post-immunization [1–4]. Intranasally-delivered live vaccines with the ability to elicit broad mucosal as well as systemic immunity are needed; respiratory mucosal immunity is essential in restricting SARS-CoV-2 replication, which in turn would reduce SARS-CoV-2 transmission.

In response to the continuous emergence of SARS-CoV-2 variants that evade pre-existing SARS-CoV-2 immunity, intranasal vaccines expressing SARS-CoV-2 antigens from recently circulating variants are being developed as booster vaccines [5–9]. Currently, the rates of severe COVID-19 resulting in hospitalization are highest in the elderly, as well as in young children under one to two years of age (https://www.cdc.gov/covid/php/covid-net/index.html; [10]). Unlike in adults, three quarters of young children hospitalized with severe COVID do not have reports of underlying chronic conditions. A bivalent vaccine against SARS-CoV-2 and human parainfluenza virus type 3, another major pediatric respiratory pathogen, for the young pediatric population would fill an unmet need.

We previously developed parainfluenza virus-vectored intranasal vaccine candidates using bovine/human parainfluenza virus type 3 (B/HPIV3) as a pediatric vector platform. B/HPIV3 is under development as a live-attenuated vaccine to protect against HPIV3, an important respiratory pathogen for the pediatric population. B/HPIV3 is based on bovine PIV3, expressing the N, P, M and L proteins from BPIV3 for host range restriction and attenuation in humans, with the HN and F proteins replaced by those from HPIV3, representing the major HPIV3 protective antigens. B/HPIV3 expressing the F protein of human respiratory syncytial virus (HRSV) has been evaluated as a bivalent HPIV3 and HRSV vaccine and was well tolerated in HPIV3-naïve children as young as 2 months of age (Clinicaltrials.gov NCT00686075, [11, 12]).

SARS-CoV-2 is continuously evolving, resulting in new antigenic variants that circulate among the population. Among various Omicron sublineages, BA.5 became the global predominant variant in early 2022 [13]. BA.5 seemed less pathogenic than ancestral SARS-CoV-2 strains but more transmissible than earlier Omicron variants [14]. BA.5 acquired the ability to escape the immunity induced by vaccination or by previous Omicron infection [15, 16].

We previously generated recombinant B/HPIV3 that expresses the 6P prefusion-stabilized version of the SARS-CoV-2 spike (S) protein of the ancestral Wuhan-Hu-1 strain from an added gene (B/HPIV3/S-6P; [17, 18]). This vaccine candidate was designed for intranasal immunization and was immunogenic and protective against SARS-CoV-2 challenge in hamsters. In rhesus macaques, mucosal immunization with B/HPIV3/S-6P induced strong systemic and mucosal immunity, including strong S-specific CD4^+^ and CD8^+^ T cell responses, and was highly protective against homologous SARS-CoV-2 challenge [18]. This vaccine is currently being evaluated in a phase I clinical study in healthy adults, as part of a step-down evaluation of safety and immunogenicity in young children (Clinicaltrials.gov NCT06026514). Recently, we evaluated the immunogenicity and protective efficacy of B/HPIV3 expressing S from the Delta or Omicron/BA.1 variants in the hamster model [19]. These vaccine candidates also induced robust immunity and were highly protective against homologous or heterologous SARS-CoV-2 challenge. However, the immune response to antigen-matched S antigens and the protective efficacy against challenge with matched SARS-CoV-2 variants exceeded that against antigenically distant versions [19], warranting further improvement of the immunization strategy. In past hamster or nonhuman primate studies of B/HPIV3-vectored COVID vaccine candidates, SARS-CoV-2 challenge was performed no later than 5 weeks after intranasal immunization [17–20]. One of the salient problems of mucosal immunity is that mucosal antibodies wane relatively quickly. In preclinical animal studies, live intranasal vaccines are effective in protecting the mucosal site of SARS-CoV-2 infection and replication, but the duration of protection elicited by mucosal vaccines warrants further characterization. In addition, since the SARS-CoV-2 S protein continues to evolve antigenically, S vaccine antigens require updating to match current strains. In the present study, we explored ways to increase the antigenic breadth of vaccines by co-administering a combination of B/HPIV3 vaccine candidates encoding antigenically distant S antigens. We performed a long-term immunization/challenge study in hamsters using two B/HPIV3 vectors expressing (i) the prefusion stabilized S antigen of the ancestral isolate of SARS-CoV-2 and (ii) a prefusion stabilized version of the SARS-CoV-2 BA.5/Omicron S protein, matching the variant that was circulating when this study was designed. To get a better take on the durability of protection elicited by these intranasal B/HPIV3 vectors, the hamster study included a SARS-CoV-2 challenge 5 months after intranasal immunization.

## Results

### Generation of B/HPIV3/S-BA.5-2P

B/HPIV3 is a recombinant live-attenuated intranasal HPIV3 vaccine candidate based on bovine PIV3, with the BPIV3 HN and F genes replaced by those from HPIV3. In previous studies, we used B/HPIV3 as a live vector vaccine candidate expressing the SARS-CoV-2 S antigen from an added gene; the generation and amplification of B/HPIV3/S-6P expressing the prefusion-stabilized S of the ancestral Wuhan-Hu-1 strain was previously described [17].

For the present study, we generated the version B/HPIV3/S-BA.5-2P, which expresses the full-length S protein derived from SARS-CoV-2 BA.5/Omicron (Fig. 1A). To generate the S-BA.5-2P ORF, the sequence of the S antigen was codon-optimized for increased expression in human cells, and two proline substitutions were introduced at aa positions 986 and 987 to stabilize the protein in the prefusion form [21]. In addition, the S1/S2 polybasic furin cleavage motif “RRAR” was ablated by “GSAS” substitution [21]. The resulting S-BA.5-2P ORF, framed by BPIV3 gene start and gene end signals, was inserted between the BPIV3 N and P genes into a full-length cDNA plasmid encoding the antigenome of B/HPIV3 [22]. B/HPIV3/S-BA.5-2P was recovered from cDNA by reverse genetics as previously described [23] and passaged once on Vero cells to generate a P2 working stock. The sequence of the B/HPIV3/S-BA.5-2P virus working stock was confirmed by Sanger sequencing.

**Figure 1.**
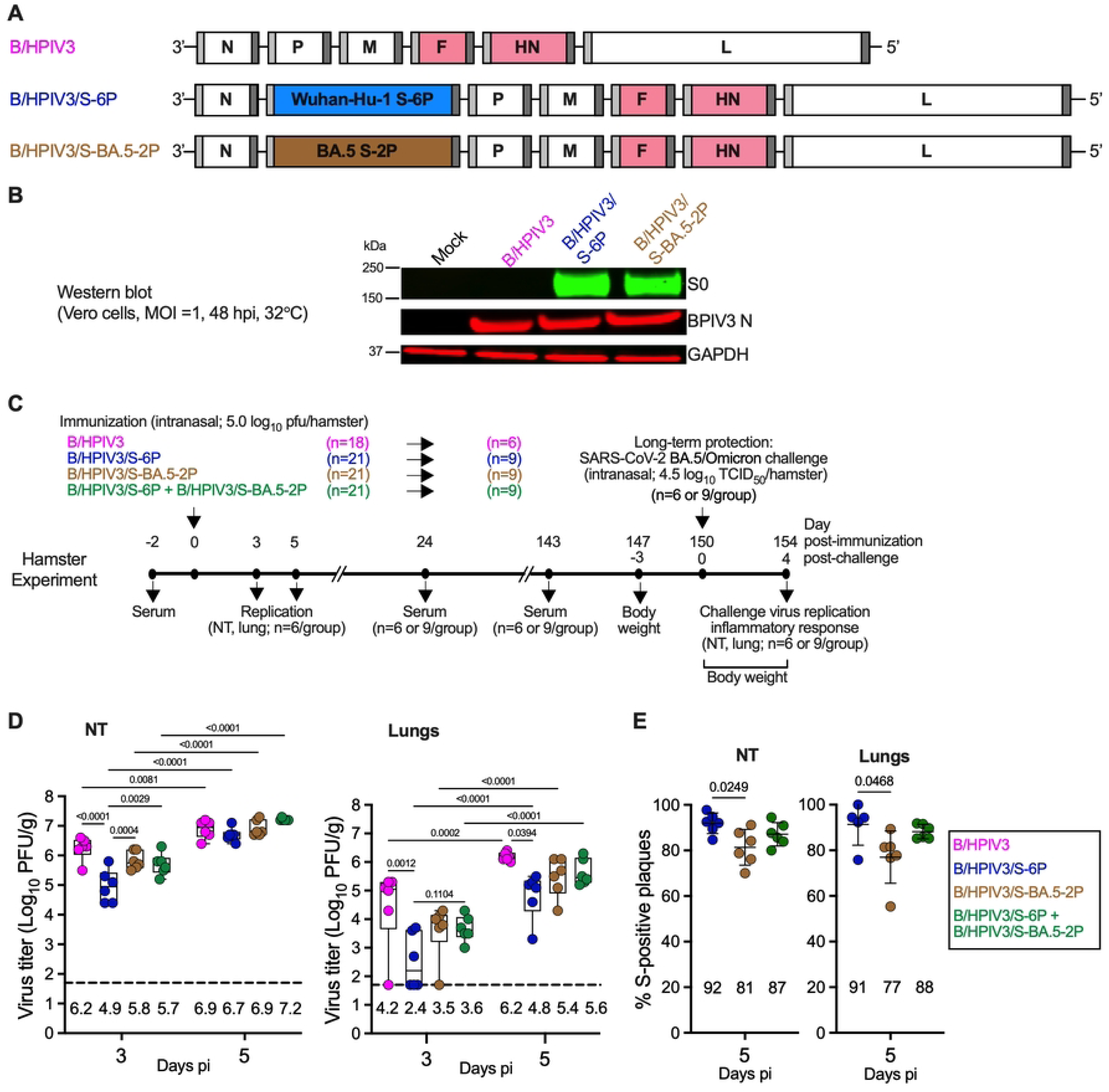
Timeline of the hamster experiment and replication of B/HPIV3 and the B/HPIV3 S-expressing vectors. (A) Genome map of B/HPIV3 empty vector and B/HPIV3 vectors encoding SARS-CoV-2 spike (S). Sequences corresponding to S (aa 1-1,273) of the ancestral Wuhan-Hu-1 or BA.5/Omicron strains were codon-optimized for expression in humans; the furin cleavage site “RRAR” was removed and replaced by “GSAS” [21], and six (S-6P) or two (S-2P) proline substitutions, respectively, were introduced to stabilize the expressed S in the prefusion form [21, 49]. S from the ancestral Wuhan-Hu-1 or BA.5/Omicron strain was inserted into B/HPIV3 to generate the previously described B/HPIV3/S-6P vector (blue) [18, 20] or B/HPIV3/S-BA.5-2P (brown). BPIV3 N, P, M and L genes are in white and HPIV3 F and HN genes in pink. The S open reading frame (ORF) was inserted between the BPIV3 N and P gene as described previously [18, 20]. Each gene is flanked by a PIV3 gene start and a gene end signal sequence (light and dark grey, respectively). (B) Subconfluent monolayers of Vero cells in 6-well plates were mock-infected or infected with the indicated virus at an MOI of 1 PFU/cell and incubated at 32°C. At 48 hpi, cell lysates were harvested and analyzed by western blotting under denaturing and reducing conditions. The SARS-CoV-2 S protein (top lane), the BPIV3 N protein (middle lane) and glyceraldehyde-3-phosphate dehydrogenase (GAPDH; included as loading control, bottom lane) were detected using primary antibodies (see Methods) followed by immunostaining with infrared fluorophore labeled secondary antibodies and infrared imaging. Images were acquired and analyzed using Image Studio software (LiCor). The approximate molecular weight in kDa is shown on the left. (C) Timeline of long-term hamster experiment. Two days before immunization, serum was collected from 81 five-to six-week-old golden Syrian hamsters. Then, three groups of hamsters (n=21 per group) were immunized intranasally with 5.0 log_10_ PFU of B/HPIV3/S-6P, B/HPIV3/S-BA.5-2P or a mixture containing 4.7 log_10_ PFU of B/HPIV3/S-6P and 4.7 log_10_ PFU of B/HPIV3/S-BA.5-2P (5.0 log_10_ PFU total). An additional group of 18 hamsters was inoculated intranasally with 5.0 log_10_ PFU of the empty vector control B/HPIV3. On days 3 and 5 post-immunization (pi), six hamsters per group were euthanized and their NTs and lungs were harvested, homogenized and aliquoted for further analysis. On days 24 and 143 pi (about 5 months pi), serum was collected from the remaining hamsters (6 or 9 hamsters per group). Hamsters were transferred to BSL3 and on day 150 pi, all remaining hamsters were challenged intranasally with 4.5 log_10_ TCID_50_/hamster of SARS-CoV-2 BA.5/Omicron. The body weight of each hamster was monitored from day 0 to day 4 post-challenge (pc). On day 4 pc, all hamsters were euthanized, and their NTs and lungs were harvested, homogenized and aliquoted for further analysis. (D) Virus titers in NTs (left panel) and lungs (right panel) on days 3 and 5 pi determined by dual-staining immunoplaque assay. Two-way ANOVA with Sidak post-test; exact p values are indicated for levels of significance p<0.05. (E) Stability of S expression by the B/HPIV3 vectors in NTs (left panel) and lungs (right panel) at day 5 pi, determined by dual-staining immunoplaque assay. Data are expressed as percent of B/HPIV3 plaques positive for S expression. N = 6 hamsters per group except the B/HPIV3/S-6P group in the lungs (n = 5) as one animal exhibit low level of virus replication. One-way ANOVA with Sidak post-test; exact p values are indicated for levels of significance p<0.05. (D and E) Medians (lines), min and max values (whiskers), 25^th^ to 75^th^ quartiles (boxes), and individual values are shown. The limit of detection is 1.7 log_10_ PFU/g (dotted line). The geometric mean titer (GMT) for each group is indicated.

### B/HPIV3/S-BA.5-2P efficiently expressed SARS-CoV-2 BA.5 S *in vitro*

We evaluated the expression of SARS-CoV-2 BA.5 S by the B/HPIV3 vector in Vero cells. Monolayers of Vero cells in 6-well plates were mock-infected or infected with B/HPIV3, B/HPIV3/S-6P or B/HPIV3/S-BA.5-2P at a multiplicity of infection (MOI) of 1 plaque forming unit (PFU)/cell and incubated at 32°C (Fig. 1B). At 48 h post-infection (pi), cell lysates were harvested, and S expression was evaluated by western blot. Expression of the BPIV3 N protein was also evaluated in parallel, and GAPDH is shown as loading control. Similarly to B/HPIV3/S-6P, B/HPIV3/S-BA.5-2P appears to efficiently express the full-length uncleaved form of SARS-CoV-2 S (S0, Fig. 1B). We also evaluated the stability of S expression by dual-staining immunoplaque assay and found that 93.3 % (±2.5%) of B/HPIV3/S-BA.5-2P plaques were positive for SARS-CoV-2 S expression. This was comparable to the S expression rates of B/HPIV3/S-6P (96.7% ±1.7% of S-positive plaques [17]). Thus, S expression by B/HPIV3 vectors expressing S-BA.5-2P or S-6P was stable.

### Replication of B/HPIV3/S-expressing vectors in hamsters

We designed a study in golden Syrian hamsters with three objectives. (i) as in our previous studies, we evaluated the replication of the B/HPIV3 vector and stability of expression of the S antigen from an added gene during *in-vivo* replication. In addition, we focused on (ii) the evaluation of the long-term protective efficacy and (iii) the effect on antigenic breadth of co-administration of B/HPIV3 expressing S proteins from distinct antigenic subgroups. Fig. 1C shows the design and the timeline of the experiment. On day 0, 5-to 6-week-old hamsters were immunized intranasally with 5 log_10_ PFU of the empty vector control B/HPIV3 (group 1, n=18). Groups 2 and 3 (n=21 per group) received 5 log_10_ PFU of B/HPIV3/S-6P or B/HPIV3/S-BA.5-2P, respectively, and group 4 (n=21) received a mixture of 4.7 log_10_ PFU of B/HPIV3/S-6P and 4.7 log_10_ PFU of B/HPIV3/S-BA.5-2P (co-administered for a total dose of 5 log_10_ PFU of vaccine) to compare the antigenic breadth of the immune response to individual candidate versus that after co-administration (81 hamsters in total; Fig. 1C).

As expected and previously described [17, 19, 20, 24], no clinical signs or symptoms of respiratory illness were observed in any of the hamsters after inoculation with either the empty B/HPIV3 vector or one or both of the S-expressing B/HPIV3 vectors. To evaluate vaccine replication, on days 3 and 5 post-immunization (pi), six hamsters per group per day were euthanized and their nasal turbinates (NTs) and lungs were harvested, homogenized and aliquoted. As expected, lungs appeared normal without gross pathology on both days. Titers of the empty vector or vaccine viruses in NT and lung homogenates were determined by dual-staining immunoplaque assay.

B/HPIV3 replicated efficiently in NTs, as typically observed, reaching a maximum geometric mean titer (GMT) of 6.9 log_10_ PFU/g of tissue on day 5 pi (Fig. 1D, left panel). Replication of all vectors significantly increased from day 3 to day 5 pi. With the exception of B/HPIV3/S-6P which was detected at significantly lower titers in NTs on day 3 pi (p<0.0001 compared to B/HPIV3 empty vector control), GMTs in NT of the S-expressing B/HPIV3 vectors on days 3 and 5 pi were overall comparable to that of B/HPIV3.

In lungs, B/HPIV3 titers significantly increased from day 3 to day 5 pi, reaching a GMT of 6.2 log_10_ PFU/g, as typically observed (Fig. 1D, right panel). Similarly, titers of the S-expressing B/HPIV3 vectors significantly increased in lungs from day 3 to day 5 pi. However, on day 5 pi, titers of the S-expressing vectors were between 4- and 25-fold lower than those of B/HPIV3, with B/HPIV3/S-6P titers being lowest (p= 0.0294 compared to B/HPIV3).

We also evaluated the stability of S expression by the B/HPIV3 vectors in NTs and lungs at day 5 pi by dual-staining immunoplaque assay (Fig. 1E). Between 81-92% and 77-91% of the plaques from NT and lungs, respectively, expressed the S antigen, indicating robust stability of S expression by the B/HPIV3 vectors in the upper and lower airways. Expression of S of the SARS-CoV-2 Omicron/BA.5 strain by B/HPIV3 however appeared slightly but significantly less stable in hamsters than that of the S protein derived from the ancestral Wuhan-Hu-1 strain.

### Short- and long-term serum SARS-CoV-2 S antibody responses in immunized hamsters

To evaluate the magnitude, breadth and long-term durability of the antibody responses to the S antigens expressed by the different S-expressing B/HPIV3 vectors, sera were collected before immunization (day -2) and on days 24 and 143 pi (corresponding to 3.5 weeks and about 5 months after immunization) from the remaining hamsters (n=6 for the B/HPIV3 group and n=9 for the other groups, see Fig. 1C). Serum anti-S and anti-receptor binding domain (RBD) IgG and anti-S IgA ELISA titers on days -2, 24 and 143 pi were determined using purified S-6P or RBD derived from Wuhan-Hu-1 (Fig. 2A and B, respectively) or purified S-2P derived from BA.5 (Fig. 2C) as antigens.

**Figure 2.**
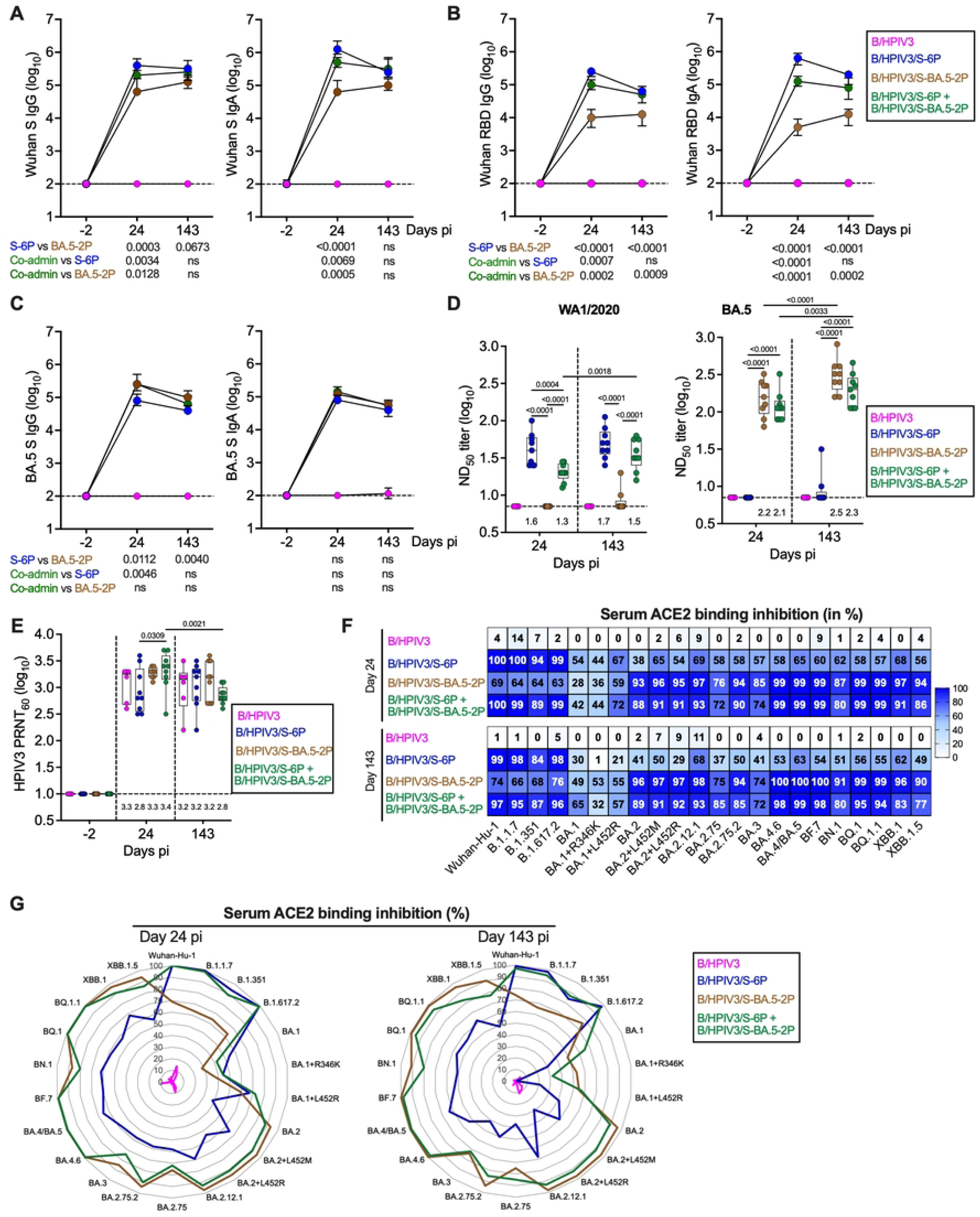
Magnitude, breadth and durability of the serum antibody response in immunized hamsters. Sera collected two days before immunization and on days 24 and 143 pi from matched hamsters (n=6 for B/HPIV3 group; n=9 for other groups, see Fig. 1C for timeline) were subjected to IgG/IgA dual ELISA (A-C), neutralization assays (D-E) and an ACE2 binding inhibition assay (F-G). (A-C) Serum IgG/IgA ELISA titers to S (6P version, A) or RBD (B) of the ancestral SARS-CoV-2 Wuhan-Hu-1 or to S (2P version) of the BA.5/Omicron variant (C). The limit of detection (dotted line) is 2 log_10_. Medians and interquartile ranges are shown. Statistical differences are shown underneath each time point (mixed-effects analysis with Sidak multiple comparison test; exact p values are indicated for levels of significance p<0.05). (D-E) Sera were also analyzed for neutralizing activities against live SARS-CoV-2 WA1/2020 (D, left panel), BA.5/Omicron (D, right panel) or HPIV3 (E). Neutralizing activities are shown as ND_50_ (D) or as 60% plaque reduction neutralization test titers (PRNT_60_, E) in log_10_. Medians (lines), min and max values (whiskers), 25^th^ to 75^th^ quartiles (boxes), and individual values are shown. The limit of detection is 0.85 (D) or 1 (E) log_10_. Median ND_50_ (D) or PRNT_60_ (E) is indicated above x axes. Mixed-effects analysis with Sidak post-test; exact p values are indicated for levels of significance p<0.05. For clarity, statistical differences against B/HPIV3 negative control (D) or against day -2 pi PRNT_60_ (E) are not shown. N=6 for B/HPIV3 group; n=9 for other groups, with the exception of n = 8 for B/HPIV3/S-BA.5-2P at 24 dpi in HPIV3 neutralization assay due to limited volume of serum for one hamster. (F-G) Binding inhibition of ACE2 protein to SARS-CoV-2 S proteins represented as a heatmap (F) or radar plots (G). Sera collected on days 24 and 143 pi were evaluated for their ability to inhibit the binding between human soluble ACE2 and the S proteins from the indicated SARS-CoV-2 strains and shown as median percentage inhibition (n=6 for B/HPIV3 group; n=9 for other groups).

Immunization with B/HPIV3/S-6P, B/HPIV3/S-BA.5-2P, or co-administration of the two S-expressing viruses induced robust anti-S and anti-RBD serum IgG and IgA titers (Fig. 2A-C). On day 24 pi, serum IgG and IgA titers to the ancestral Wuhan-Hu-1 S and RBD in the B/HPIV3/S-6P groups significantly exceeded those in the B/HPIV3/S-BA.5-2P group (Fig. 2A and B). On the other hand, immunization with B/HPIV3/S-BA.5-2P induced significantly higher IgG titers to BA.5/Omicron S antigen than B/HPIV3/S-6P (Fig. 2C). Co-administration of the two S-expressing vectors induced higher IgG and IgA ELISA titers on day 24 against ancestral S and RBD than B/HPIV3/S-BA.5-2P alone, as well as higher IgG titers to S-BA.5 than B/HPIV3/S-6P alone, suggesting increased antibody breadth (Fig. 2C). The anti-S and anti-RBD IgG and IgA ELISA titers induced by each immunization regimen remained high up to day 143 pi (about 5 months pi). As expected, no anti-S or anti-RBD antibodies were detected on day -2 pi or in hamsters immunized with the empty B/HPIV3 vector control.

We also evaluated the neutralizing activity of the sera collected on days 24 and 143 pi against live SARS-CoV-2 WA1/2020 or BA.5/Omicron (BSL3) (Fig. 2D); the neutralizing antibodies to HPIV3 were also evaluated (BSL2) (Fig. 2E). B/HPIV3/S-6P induced robust levels of serum neutralizing antibodies against the matched WA1/2020 strain (median 50% neutralizing dose (ND_50_) titer of 1.6 log_10_ on day 24 pi; Fig. 2D, left panel), with a slight increase by 5 months after immunization. However, sera from these animals did not neutralize BA.5/Omicron (Fig. 2D, right panel). On the other hand, sera from the B/HPIV3/S-BA.5-2P-immunized hamsters did not neutralize the WA1/2020 strain (Fig. 2D, left panel) but induced high levels of neutralizing antibodies against the matched BA.5/Omicron virus (median ND_50_ titer of 2.2 log_10_ on day 24 pi; Fig. 2D, right panel). Five months after immunization, the homologous BA.5 neutralizing titers had further increased (p<0.0001 between day 24 and day 143 pi). Co-administration of the two S-expressing vectors induced robust titers of serum neutralizing antibodies against both WA1/2020 and BA.5/Omicron strains (median ND_50_ titers of 1.3 to 2.1 log_10_ on day 24 pi, respectively, Fig 2D left and right panel), confirming the increased breadth of serum antibody responses in response to co-administration. Five months after immunization, these titers had slightly but significantly increased (p=0.0018 and p=0.0033 for ND_50_ titers against WA1/2020 and BA.5/Omicron, respectively). Except for the WA1-2020 neutralizing titers elicited by B/HPIV3/S-6P alone on day 24, the antigen-matched neutralizing titers induced by the single vaccine candidates did not significantly exceed those in the co-administration groups. As expected, serum from hamsters immunized with the empty vector control B/HPIV3 did not neutralize either the WA1/2020 or BA.5/Omicron strains. In addition, sera collected on day 24 pi from all immunized hamsters efficiently neutralized HPIV3 (GMTs between 2.9-3.3 log_10_, Fig. 2E). Five months after immunization, HPIV3 neutralizing titers remained high in all groups with nevertheless a modest but significant reduction in the co-administration group (Fig. 2E).

The breadth of the SARS-CoV-2 antibody response was further evaluated in an angiotensin-converting enzyme 2 (ACE2) binding inhibition assay (Fig. 2F-G). This assay evaluates the ability of serum antibodies to inhibit the binding of soluble human ACE2 receptor to S antigens of different SARS-CoV-2 strains. Sera collected on day 24 pi from B/HPIV3/S-6P-immunized hamsters efficiently inhibited binding of ACE2 to S from Wuhan-Hu-1 and early variants of concern (B.1.1.7/Alpha, B.1.351/Beta and B.1.617.2/Delta strains; 94 to 100% inhibition; Fig. 2F, top panel, and 2G left panel). However, serum inhibition of ACE2 binding to S from BA.1, BA.2, BA.3, BA.4 and derivatives as well as more recently circulating variants (BF.7, BN.1, BQ.1, XBB.1 and derivatives) was lower (38 to 69% inhibition). On the other hand, sera from B/HPIV3/S-BA.5-2P-immunized hamsters efficiently inhibited the binding of ACE2 to S from BA.2, BA.3, BA.4, BA.5 and derivatives as well as from more recently circulating variants (76 to 99% inhibition). However, sera from these hamsters exhibited reduced ACE2 binding inhibition to S from early variants of concern or BA.1/Omicron and derivatives (28 to 69% inhibition). Interestingly, sera from hamsters immunized with the mixture of both B/HPIV3-S expressing vectors showed the combined binding inhibition profile of sera from animals immunized with each of the individual S-expressing vectors, confirming the increased antibody breadth of the immune response. Sera from these hamsters highly efficiently inhibited the binding of ACE2 to S from almost all variants (72 to 100% inhibition), with the exception of BA.1 and derivatives, for which the inhibition was lower (42 to 72% inhibition). Comparable results were obtained using sera collected on day 143 pi (Fig. 2F bottom panel and 2G right panel), suggesting that the breadth of the antibody response was maintained over almost 5 months after immunization. As expected, sera from the B/HPIV3 empty vector control group did not inhibit binding of soluble ACE2 to S.

### Long-term protection of immunized hamsters against weight loss and inflammatory responses following BA.5/Omicron challenge

We next evaluated the long-term protection induced by the various immunization regimens against challenge with SARS-CoV-2 BA.5/Omicron. All remaining hamsters (n=6 for the B/HPIV3 control group and n=9 for the other groups) were challenged intranasally on day 150 pi (5 months pi) with 4.5 log_10_ 50% tissue culture infection dose (TCID_50_) of SARS-CoV-2 BA.5/Omicron (see Fig. 1C for timeline of the experiment). Although the BA.5/Omicron strain induces only modest weight loss in the hamster model [25], animals were monitored for weight change from day 150 pi (the challenge day) through day 154 pi (day 4 post-challenge, pc). On day 4 pc, all animals were necropsied, and NT and lungs were harvested, homogenized, aliquoted and stored before further analysis.

Hamsters immunized with the B/HPIV3 empty vector control experienced a modest weight loss after challenge (median weight loss of 1.6%, with two hamsters showing 6.9% and 4.3% weight loss, respectively, Fig. 3A). In the three other groups of hamsters immunized with the S-expressing vectors, weight remained overall steady from day 0 to day 2 pc and then increased on days 3 and 4 pc. This suggested that hamsters immunized with the S-expressing B/HPIV3 vectors were protected from weight loss.

**Figure 3.**
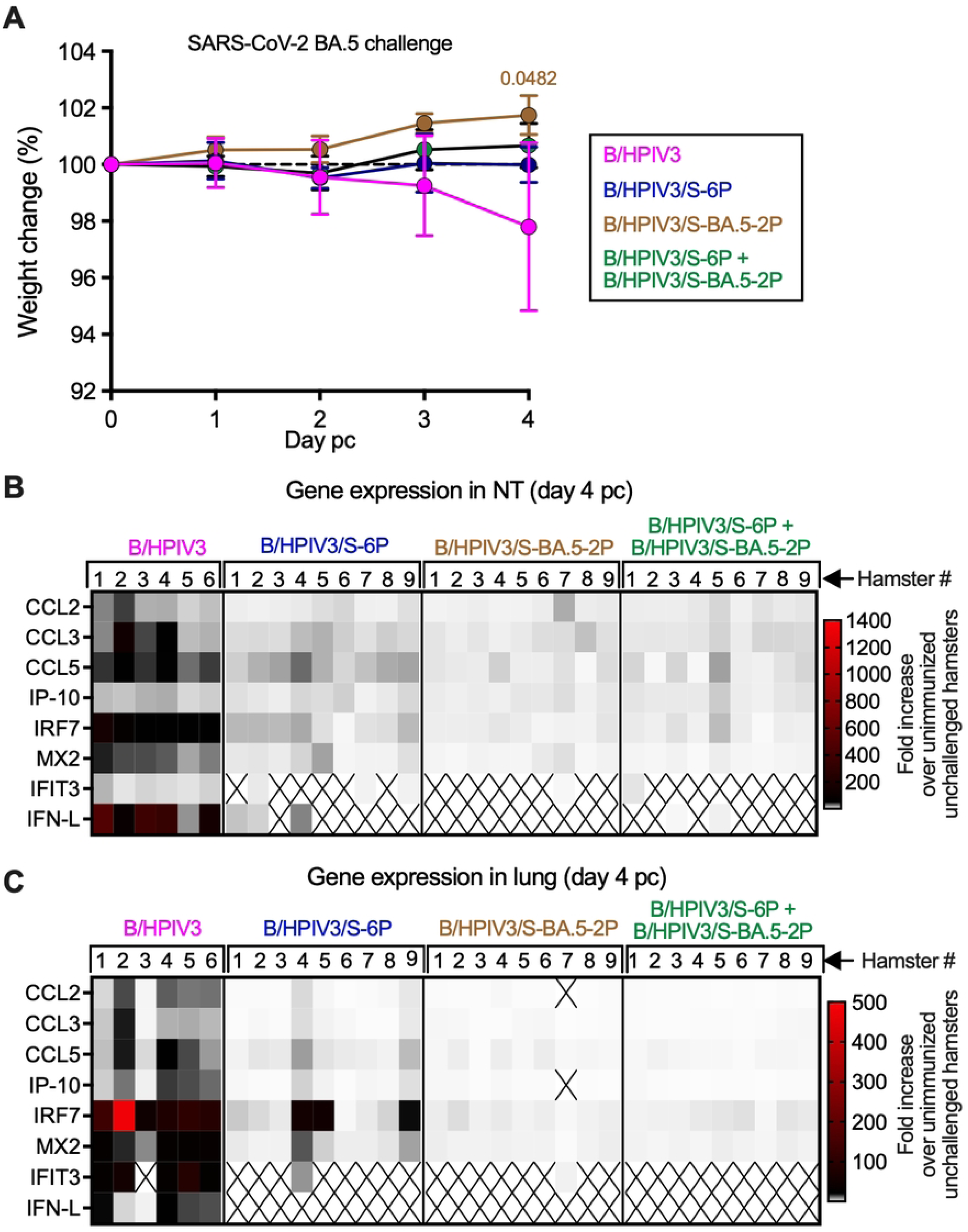
Body weight changes and inflammatory responses in upper and lower airways of immunized hamsters challenged with SARS-CoV-2 BA.5/Omicron. On day 150 pi, remaining hamsters (n=6 for B/HPIV3 group; n=9 for other groups) were challenged intranasally with 4.5 log_10_ TCID_50_ per animal of BA.5/Omicron. Body weights were monitored daily from day 0 to 4 post-challenge (pc). On day 4 pc, all hamsters were euthanized, and NTs and lungs were collected, homogenized and aliquoted for further analysis (see Fig. 1B for timeline). (A) Body weight changes from day 0 to day 4 pc shown as mean percent body weight with standard deviation relative to the day 0 weight. Mixed-effects analysis with Dunnett post-test; exact p value is indicated for levels of significance p<0.05. (B-C) Expression of inflammatory/antiviral genes in NTs (B) and lungs (C) on day 4 pc. Total RNA was extracted from NT and lung homogenates followed by cDNA synthesis for TaqMan assays. Results were analyzed using the ΔΔCt method and normalized to β actin. Data are represented as fold increase over the mean level of expression in NTs or lungs of six or five unimmunized unchallenged hamsters, respectively, harvested from previously published studies [17, 20]. As the expression of IFIT3 and IFN-L was not detected in any of the control hamsters, their level of expression was determined relative to the immunized/challenged hamster with the lowest detectable level of expression (see methods). The expression of IFIT3 in NT and lungs was determined relative to hamster #7 of the B/HPIV3/S-BA.5-2P group. The expression of IFN-L in NT was determined relative to hamster #3 of the B/HPIV3/S-6P+B/HPIV3/S-BA.5-2P group and, in the lungs, relative to hamster #3 of the B/HPIV3 group.

Inflammatory responses in the upper and lower respiratory tract were also evaluated following challenge. Total RNA was extracted from NT and lung homogenates collected on day 4 pc, and the level of expression of a subset of inflammatory/antiviral genes (CCL2, CCL3, CCL5, IFIT3, IFN-L, IP-10, IRF7, MX2) was determined by RT-qPCR (Fig. 3B and C). Data are represented as fold increase over the mean level of expression in NTs or lungs of six or five unimmunized unchallenged hamsters from the same source, respectively, collected in previous studies [17, 20]. As the expression of IFIT3 and IFN-L was not detected in any of the control hamsters, their level of expression was determined relative to the immunized/challenged hamster with the lowest detectable level of expression (see methods). High levels of expression of inflammatory genes were detected in the NTs and lungs of hamsters immunized with the B/HPIV3 empty vector control and challenged with BA.5/Omicron (Fig. 3B and C, left panels). A mild but detectable inflammatory response was detected in the NTs of hamsters immunized with B/HPIV3/S-6P after SARS-CoV-2 BA.5 challenge (Fig. 3B). However, the inflammatory response appeared minimal in the NT of hamsters immunized intranasally with B/HPIV3/S-BA.5-2P or the combination of the two S-expressing vectors (Fig. 3B).

In the lungs (Fig. 3C), with the exception of two animals in the group of hamsters immunized with the B/HPIV3/S-6P vector that exhibited moderate inflammatory responses (hamsters #4 and #9), all hamsters immunized with the S-expressing vectors appeared protected from inflammatory responses following challenge with BA.5/Omicron (Fig. 3C).

### Long-term protection of immunized hamsters against BA.5/Omicron challenge virus replication

SARS-CoV-2 BA.5/Omicron challenge virus titers were determined in homogenates of NT and lung tissues collected on day 4 pc (Fig. 4A and B). As expected, BA.5/Omicron replicated to high level in NTs and lungs of hamsters immunized with the empty B/HPIV3 vector control (5.7 and 5.5 log_10_ TCID_50_/g of tissue in the NTs and lungs, respectively). BA.5/Omicron challenge virus was also detected in the NTs of hamsters immunized with B/HPIV3/S-6P, but titers were about 400-fold lower compared to B/HPIV3 empty-vector immunized hamsters. No challenge virus was detectable in the NTs of hamsters immunized with the vaccine-matched B/HPIV3/S-BA.5-2P vector alone nor with the combination of the two S-expressing B/HPIV3 vectors (Fig. 4A). None of the hamsters immunized with the S-expressing vectors had challenge virus detectable in lung tissues, indicating that they were fully protected against BA.5/Omicron replication in the lungs (Fig. 4B).

**Figure 4.**
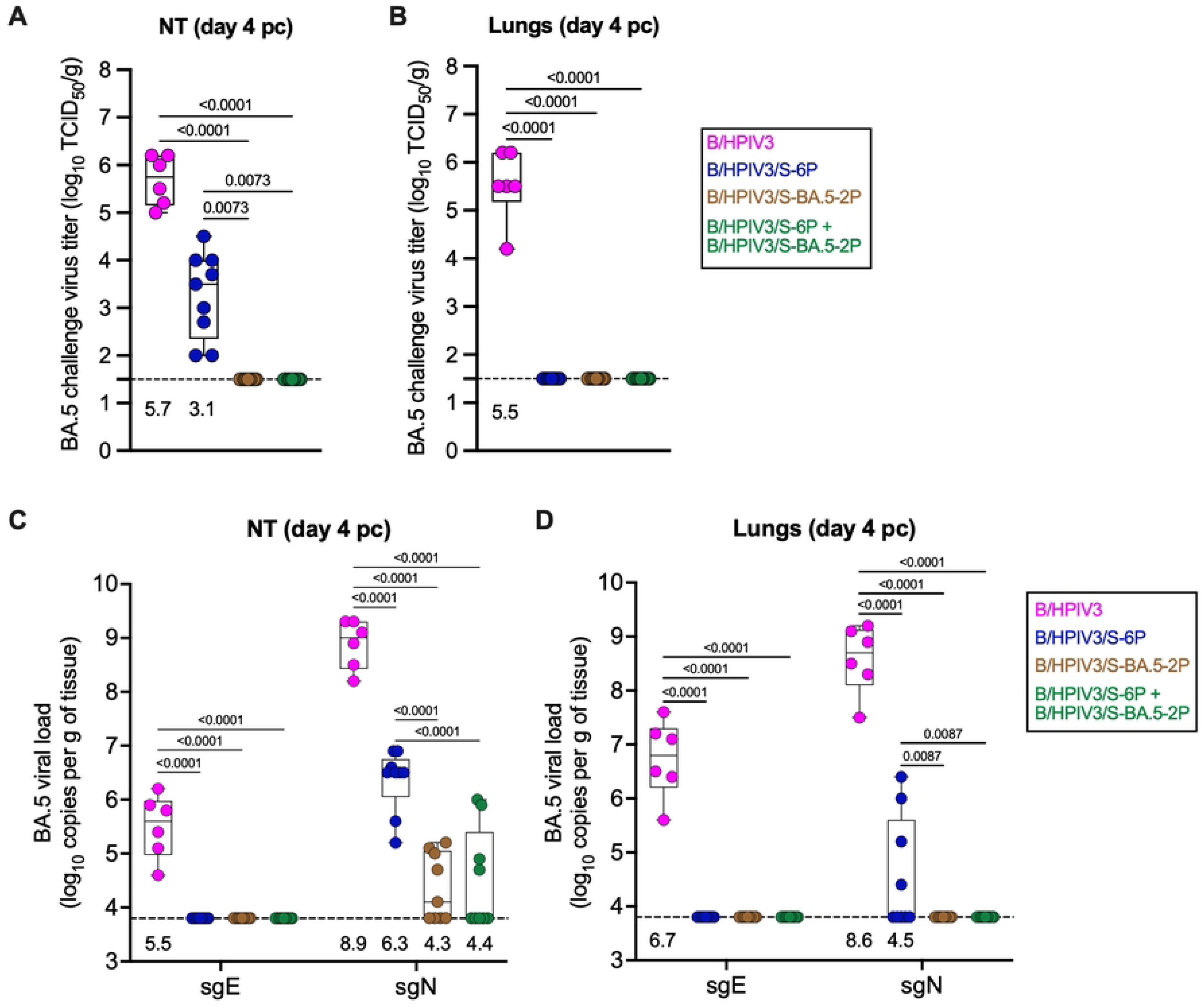
Protection of immunized hamsters against SARS-CoV-2 BA.5/Omicron challenge virus replication. Replication of the challenge virus was evaluated from NT and lung homogenates collected on day 4 pc. Quantification of SARS-CoV-2 BA.5/Omicron viral genomes was also performed using total RNA isolated from additional aliquots of NT and lung homogenates (n=6 for B/HPIV3 group; n=9 for other groups). **(A-B) Titers of SARS-CoV-2 challenge virus BA.5 were determined from NT (A) and lung** homogenates (B) and expressed in log_10_ TCID_50_ per g of tissue. The limit of detection is 1.5 log_10_ TCID_50_ per g. (C-D) SARS-CoV-2 viral loads in NTs (C) and lungs (D) were determined from total RNA extracted from tissue homogenates by using TaqMan assays to detect sgE and sgN mRNA. The limit of detection is 3.8 log_10_ copies per g of tissue. Medians (lines), min and max values (whiskers), 25^th^ to 75^th^ quartiles (boxes), and individual values are shown. GMTs are also indicated above x axes. Two-way ANOVA with Sidak post-test; exact p values are indicated for levels of significance p<0.05.

Challenge virus genome replication was also quantified in both NTs and lungs by RT-qPCR (Fig. 4C and D). To do so, total RNA was extracted from tissue homogenates, and SARS-CoV-2 subgenomic E (sgE) and N (sgN) mRNA were quantified as an indicator of active virus replication. In NTs from hamsters immunized with the B/HPIV3 empty vector control, high levels of sgE and sgN mRNAs were detected (GMT of 5.5 and 8.9 log_10_ copies/g, respectively; Fig. 4C). In NTs from hamsters previously immunized with the S-expressing vectors, no sgE mRNA and significantly reduced levels of sgN mRNA were detected after challenge (Fig. 4C). More specifically, hamsters immunized with B/HPIV3/S-6P alone exhibited a 400-fold lower level of sgN mRNA compared to B/HPIV3-immunized hamsters (GMT of 6.3 log_10_ copies/g). Hamsters immunized with the vaccine-matched B/HPIV3/S-BA.5-2P or the combination of the two S-expressing B/HPIV3 vectors exhibited a 500-fold lower level of sgN mRNA. In the lungs, high levels of sgE and sgN mRNA were detected in hamsters immunized with the B/HPIV3 empty-vector control (GMT of 6.7 and 8.6 log_10_ copies/g, respectively; Fig. 4D). Low levels of sgN mRNA were detected only in a subset of hamsters immunized with B/HPIV3/S-6P. No sgE or sgN mRNA was detected in B/HPIV3/S-BA.5-2P immunized hamsters or hamsters that had received the combination of vaccine candidates (Fig. 4D).

## Discussion

SARS-CoV-2 continues to evolve antigenically, and symptomatic reinfections with newly circulating variants are common. The upper respiratory tract is the primary site of SARS-CoV-2 infection and replication, and broad and durable mucosal immunity is critical in limiting infection and spread. Based on antigenic cartography studies, the ancestral Wuhan-Hu-1 SARS-CoV-2 strain is antigenically distant from a cluster of variants centering around BA.4/BA.5 Omicron lineages. We are developing live-attenuated B/HPIV3-vectored vaccines expressing SARS-CoV-2 S antigen as bivalent intranasal vaccines to protect children against HPIV3 and COVID-19 [17–19]. The safety, immunogenicity and protective efficacy of B/HPIV3-S-expressing vaccines have been described in our previous preclinical studies in hamsters and rhesus macaques [17, 18]. Based on these data, B/HPIV3/S-6P expressing the ancestral S is currently being evaluated in a phase I clinical trial (Clinicaltrials.gov NCT06026514). In our most recent preclinical study [19], we compared the immunogenicity and breadth of protection of the B/HPIV3-expressed S protein of the ancestral, B.1.617.2/Delta, and B.1.1.529/Omicron variants in hamsters. All three intranasal SARS-CoV-2 vaccine candidates induced cross-protective SARS-CoV-2 immunity, albeit with limited antigenic breadth, similar to that of injectable SARS-CoV-2 vaccines. It became clear that vaccines with the ability to elicit broad S-specific antibody responses to a broad range of antigenically-distinct SARS-CoV-2 variants are needed. Antigenic cartography places the S antigen of ancestral SARS-CoV-2 in a distant relationship to BA.5/Omicron (and more recent) variants [26–30]. In the present study, we used versions of B/HPIV3 expressing the ancestral or the BA.5 S antigen, alone or in combination, to explore ways to increase the antigenic breadth of these vaccine candidates. Administered individually, both candidates induced a strong antigen-matched S-specific response, as expected based on previous studies for the ancestral antigen, and as predicted for the new BA.5/Omicron S-expressing candidate. We further evaluated the breadth of serum antibody responses by ELISA, neutralization assay, and the highly sensitive ACE2 binding inhibition assay. The ACE2 inhibition assays showed that intranasal immunization of hamsters using a mixture of B/HPIV3/S-6P expressing S from the ancestral SARS-CoV-2 strain and B/HPIV3/S-BA.5-2P induced a broad antibody response to pre-Omicron and Omicron variants, as well as XBB strains, combining the breadth of the antibody response induced by each B/HPIV3-S expressing vector individually. We found no evidence of interference or restriction between the two vaccine candidates when administered together. A previous study evaluating a bivalent injectable vaccine based on adenovirus vectors expressing S of the ancestral and BA.5/Omicron strains also exhibited a broad neutralizing antibody response, but XBB strains were poorly neutralized [31]; an injectable nanoparticle-based vaccine displaying the RBDs of the ancestral and Omicron/BA.5 strains induced a broad antibody response against recently circulating isolates including XBB variants [32], similarly to the antibody breadth elicited by the intranasal vaccines in the present study. The reason for the discrepancies in the antibody breadths of antibodies elicited by these vaccines will need further investigation.

Low levels of mucosal neutralizing antibodies are typically detected after vaccination with an mRNA-based vaccine [33]. Intranasally or intratracheally delivered adenovirus-based vaccines have been evaluated in animal models as boosters to provide increased protection against Omicron strains [5–7]. These vaccines conferred broad immunity and robust protection against SARS-CoV-2 challenge, in some cases surpassing the protection provided by mRNA-based vaccines, especially in the upper airways [6, 7]. In the hamster model, mucosal immunization with an adenovirus-based vaccine or a live-attenuated SARS-CoV-2 vaccine effectively restricted replication of SARS-CoV-2 ancestral, BA.1 or BA.5/Omicron variants as well as spread and replication in the upper and lower airways of contact hamsters [8, 9]. The restrictive effects of the mucosal immunity elicited by these intranasal vaccine candidates on SARS-CoV-2 replication in the airways prevented subsequent virus transmission, while SARS-CoV-2 transmission was not restricted after intramuscular immunization with mRNA vaccine [8, 9].

B/HPIV3 efficiently and stably expressed S of BA.5/Omicron *in vitro* and to levels that were comparable to S of the ancestral, B.1.1.529/Omicron, B.1.617.2/Delta strains [17, 19]. In the present study, expression of the BA.5/Omicron S protein by B/HPIV3 in the upper and lower airways of hamsters appeared more stable than that of the B.1.1.529/Omicron S protein in our most recent study [19]. In the present study, B/HPIV3/S-BA.5-2P replicated to titers that were lower overall than that of the B/HPIV3 empty-vector control, but slightly higher than that of B/HPIV3/S-6P. We previously observed that replication of B/HPIV3 expressing S of B.1.1.529/Omicron was lower than B/HPIV3/S-6P [19]. Thus, while B/HPIV3 expressing S from different strains replicate efficiently in the respiratory tract of hamsters, the origin of S may affect vector replication and stability of S expression.

In our previous studies, we followed vaccinated hamsters for only a relatively short duration of four to five weeks, preventing the exploration of the durability of immunogenicity and protective efficacy of these vaccine candidates. The present study was designed as an extended vaccine study to evaluate the long-term immunogenicity and protective efficacy five months after immunization against challenge with BA.5/Omicron. Consistent with previous results, we detected robust serum IgG and IgA responses to the S antigen in all hamsters at 3-4 weeks pi [17, 19, 20], and the long-term durability of the serum antibody response over 5 months was excellent. Serum IgG and IgA titers to S (ancestral Wuhan-Hu-1 or BA.5/Omicron) or RBD (Wuhan-Hu-1) were generally sustained over the five-month time frame, suggesting that the level and breadth of the antibody responses induced by a single intranasal dose were stable for at least 5 months. Similarly, serum neutralizing antibody responses to vaccine-matched SARS-CoV-2 or HPIV3 were sustained over the 5 months. On the other hand, in nonhuman primates immunized parenterally with two doses of mRNA-based vaccines, neutralizing antibody responses continuously waned over 6 months [34, 35].

About five months after immunization, hamsters were challenged intranasally with a SARS-CoV-2 BA.5/Omicron isolate. In SARS-CoV-2 naïve hamsters, this lineage only induces moderate weight loss [25]. Indeed, in our study, we observed some weight loss in hamsters previously immunized with the B/HPIV3 empty vector control. Hamsters immunized with B/HPIV3/S-6P expressing the ancestral S protein did not lose weight after challenge, but based on the increase in host inflammatory gene expression and virus gene expression detected in the upper and lower airways after BA.5/Omicron challenge, we concluded that B/HPIV3/S-6P was only partially protective against BA.5/Omicron challenge. This further confirmed that antibodies induced by S antigen from the ancestral strain only induced partial cross-neutralization against BA.5/Omicron. On the other hand, hamsters immunized with B/HPIV3/S-BA.5-2P or a mixture of the two B/HPIV3/S-expressing vectors were robustly protected against challenge based on absence of weight loss, expression of host inflammatory genes in the upper and lower airways, or SARS-CoV-2 replication after challenge. Only low levels of sgN mRNA were detected in the NT. Thus, five months after intranasal immunization, the protective efficacy of B/HPIV3 expressing S from BA.5/Omicron was still sustained in hamsters. Furthermore, this suggests that the robust protection against SARS-CoV-2 replication in the upper airways might help to prevent SARS-CoV-2 transmission. While we were not able to assess efficacy against SARS-CoV-2 transmission in our present long-term study, this will be evaluated in a follow-up study.

The immunogenicity and long-term protection induced by the B/HPIV3 S-expressing vector are currently being further evaluated in the rhesus macaque model. Due to technical limitations, we were not able to measure the mucosal IgG and IgA response in hamsters and had to rely on serum IgA as a correlate of mucosal antibodies. The long-term study in macaques will allow an in-depth evaluation of the mucosal immune response. The results from this preclinical evaluation in hamsters suggest that co-administration of B/HPIV3 vectors expressing antigenically distinct S antigens should be evaluated in clinical trials to further increase the breadth of the antibody response and provide better protection against constantly evolving SARS-CoV-2 variants.

## Materials and methods

### Ethics statement

Hamster studies were approved by the Animal Care and Use Committee of the National Institute of Allergy and Infectious Diseases. The animal experiments were performed following the Guide for the Care and Use of Laboratory Animals by the NIH.

### Generation of B/HPIV3 expressing SARS-CoV-2 BA.5/Omicron antigen

The sequence encoding the full-length open reading frame of the BA.5/Omicron S protein was based on GenBank accession number QHD43416.1 [36], with D25/27, D69/70, G142D, V213G, G339D, S371F, S373P, S375F, T376A, D405N, R408S, K417N, N440K, L452R, S477N, T478K, E484A, F486V, Q493, Q498R, N501Y, Y505H, D614G, H655Y, N679K, P681H, N764K, D796Y, Q954H, and N969K mutations and 682-RRAR-to-GSAS-685 and K986P/V987P substitutions for S-2P stabilization. The codon-optimized sequence was synthesized (GenScript) and inserted into a cDNA plasmid containing the B/HPIV3 antigenome sequence as an additional gene, framed by PIV3 gene-start and gene-end signals, between the B/HPIV3 N and P genes using the same strategy as previously described [20]. Recombinant virus was recovered from cDNA by reverse genetics [23], and viral stocks were prepared on Vero cells. The sequence of recombinant B/HPIV3/S-BA.5-2P (with the exception of the most outer 5’ and 3’ sequences imposed by the primers used for amplification) was confirmed by Sanger sequencing.

### Western blotting

Viral protein expression by B/HPIV3/S-6P, B/HPIV3/S-BA.5-2P or the B/HPIV3 empty vector was evaluated as previously described [17, 19]. Briefly, Vero cells were seeded in 6-well plates and on the next day, were inoculated with P2 stocks of B/HPIV3, B/HPIV3/S-6P or B/HPIV3/S-BA.5-2P at an MOI of 1 PFU per cell. After incubation for 48 h at 32°C, cells were washed with Dulbecco’s phosphate-buffered saline (DPBS) and lysed using 300 μl per well of 1X NuPAGE LDS Sample Buffer (Thermo Fisher, NP0007). Lysates were clarified by using QIAshredders (Qiagen, 79656) and mixed with NuPAGE Sample Reducing Agent (Thermo Fisher, NP0009) before being denatured for 10 min at 90°C. The lysates were separated on NuPAGE Bis-Tris Mini Protein Gels, 4–12% (Thermo Fisher, NP0335BOX). The resolved proteins were transferred to polyvinylidene difluoride membranes included in iBlot 3 Transfer Stacks (Thermo Fisher, IB34002) using an iBlot 3 Western Blot Transfer Device (Thermo Fisher). Membranes were blocked with Intercept Blocking Buffer (LiCor) at room temperature for 1 h and incubated with primary antibodies (a goat hyperimmune serum to SARS-CoV-2 S; a rabbit polyclonal hyperimmune serum against purified B/HPIV3; a mouse monoclonal antibody to GAPDH (Sigma, G8795)) (all at 1:5,000) in Intercept Blocking Buffer containing 0.05% Tween 20 overnight at 4°C. Membranes were further incubated with infrared dye-labeled secondary antibodies from LiCor (IRDye 800CW donkey anti-goat IgG (926-32214); IRDye 680RD donkey anti-rabbit IgG (926-68073); IRDye 680RD donkey anti-mouse IgG (925-68072)) (all at 1:10,000) in Intercept Blocking Buffer containing 0.05% Tween 20 for 1 h at room temperature and scanned using an Odyssey CLx Imager (LiCor).

### SARS-CoV-2 viruses

SARS-CoV-2 USA-WA1/2020 virus was obtained through BEI Resources (cat # NR-52281) and was passaged twice on Vero-TMPRSS2 cells. The SARS-CoV-2 BA.5 isolate for hamster challenge (hCoV-19/USA/COR-22-063113/2022; lineage BA.5; Omicron variant) was obtained through BEI Resources, NIAID, NIH (#NR-58616, contributed by Dr. Richard J. Webby).

### Hamster study

Eighty-one 5-to 6-week-old male golden Syrian hamsters (*Mesocricetus auratus*) were purchased from Envigo Laboratories (Frederick, MD) and used for experiments in BSL2 and BSL3 facilities approved by the centers for disease control and prevention. The timeline of the experiment is described in Fig 1C. Two days before immunization, serum was collected from all animals, and hamsters were assigned to three groups of 21 hamsters and one group of 18 hamsters. The group of 18 hamsters was immunized intranasally under light isoflurane anesthesia with 5 log_10_ PFU of the B/HPIV3 empty vector control. The three groups of 21 hamsters each were immunized intranasally with 5 log_10_ PFU of B/HPIV3/S-6P, B/HPIV3/S-BA.5-2P, or with a combination of 4.7 log_10_ PFU of B/HPIV3/S-6P and 4.7 log_10_ PFU of B/HPIV3/S-BA.5-2P (5 log_10_ PFU total). On days 3 and 5 pi, six animals per group per day were euthanized and NTs and lungs were harvested and homogenized. Aliquots were snap-frozen in dry ice and stored at −80°C for further analysis. On days 24 and 143 pi, serum was collected from all remaining hamsters (six hamsters for the B/HPIV3 group and nine hamsters per group for the other groups, n = 33 animals total). Between days 143 and 147 pi, animals were transferred to a BSL3 facility. On day 147 pi (three days before challenge), all remaining hamsters were weighed. On day 150, all hamsters were challenged intranasally with 4.5 log_10_ TCID_50_/hamster of the SARS-CoV-2 BA.5 isolate (hCoV-19/USA/COR-22-063113/2022), and their weight was monitored daily from day 0 to 4 pc. On day 154 pi (day 4 pc), all remaining hamsters were euthanized, and their NTs and lungs were collected and homogenized. Aliquots were snap-frozen in dry ice and stored at −80°C before further analysis.

### Dual-staining immunoplaque assay

Sub-confluent Vero cells in 24-well plates were inoculated with 10-fold serially diluted NT or lung homogenates and were incubated for 2 h at 32°C. Cells were overlaid with 1 ml per well of 0.8% methylcellulose (Sigma) dissolved in Opti-MEM (ThermoFisher) supplemented with 2% Fetal Bovine Serum, 1% L-glutamine, 2.5% penicillin-streptomycin, 0.5% amphotericin B and 0.1% gentamicin. After 7 days of incubation at 32°C, cell monolayers were fixed with ice-cold 80% methanol and blocked with Odyssey blocking buffer (LiCor). HPIV3 antigens and SARS-CoV-2 S protein were detected by dual-staining of plaques using a primary rabbit anti-HPIV3 serum at 1:5,000 [37] and a primary human anti-SARS-CoV-2 S monoclonal antibody (CR3022) at 1:2,000 [18, 38], followed by staining with a secondary IRDye 680RD donkey anti-rabbit IgG and IRDye 800CW goat anti-human IgG secondary antibodies (LiCor). Stained cells were scanned using an Odyssey CLx Imager (LiCor).

### IgG/IgA dual ELISA

Soluble versions of SARS-CoV-2 Wuhan-Hu-1 S (6P version) and RBD were purified as described previously [20]. A soluble version of SARS-CoV-2 BA.5 S (2P version) was purified as described previously [39] using a cDNA encoding the SARS-CoV-2 BA.5/Omicron S. Serum IgG/IgA titers were determined by ELISA as previously described [40]. Briefly, MaxiSorp black 96-Well immuno plates (Thermo Fisher) were coated with 75 ng per well of purified S (6P version for Wuhan-Hu-1 S; 2P version for BA.5/Omicron S) or RBD of the Wuhan-Hu-1 strain at 4°C overnight and were blocked with 5% nonfat dry milk (Nestle) in Dulbecco’s phosphate buffered saline (DPBS) (Thermo Fisher) at 4°C overnight. The following steps were all performed at room temperature and plates were washed with 1X PBS (Thermo Fisher) with 0.1% IGEPAL CA-630 (Sigma-Aldrich) between each step. First, diluted sera were prepared by 3-fold, 11-point dilutions, starting at a 1:100 dilution, using 5% milk in DPBS with 0.2% IGEPAL CA-630. Then, 100 μl of diluted sera was added to each well in duplicate plates and incubated for 1 h. Next, the plates were incubated for 1 h with a mixture of secondary antibodies: a goat anti-hamster IgG (H+L) conjugated with horseradish peroxidase (HRP, Thermo Fisher; cat# PA1-29626) diluted at 1:10,000 and a rabbit anti-hamster IgA conjugated with biotin (Brookwood Biomedical; cat# sab3002a) diluted at 1:1,000 in 5% milk in DPBS with 0.2% IGEPAL CA-630. The plates were subsequently incubated with Europium-labelled streptavidin (Perkin Elmer; cat# 1244-360) at 1:2,000 in DPBS with 0.2% IGEPAL CA-630 for 1 h. Then, plates were incubated with a 1:1 mixture of peroxide and luminol enhancer solution included in Pierce ECL Western Blotting Substrate (Thermo Fisher; cat# 32106) for 10 min and subjected to luminescence reading to quantify antigen-specific IgG using a Synergy Neo2 reader (BioTek). The plates were further incubated with DELFIA Enhancement Solution (Perkin Elmer; cat# 4001-0010) for 20 min and were subjected to fluorescence reading (excitation at 360/40; emission at 620/40) to detect antigen-specific IgA with the same reader. ELISA titers were determined as described previously [40]; (i) the average reading from duplicate wells was calculated, (ii) the average reading from blank samples was subtracted, (iii) the cut-off value was set to the blank average plus three standard deviations, and (iv) the IgG and IgA titers of each sample were determined in sequential reads by interpolating the sigmoid standard curve generated on Prism 9.0 (GraphPad Software).

### Live-virus neutralization assay

SARS-CoV-2 neutralizing antibody titers in hamster sera were determined by live-virus neutralization assays in a BSL3 facility. Heat-inactivated sera were 2-fold serially diluted in Opti-MEM and mixed with 100 TCID_50_ of SARS-CoV-2 WA1/2020 or BA.5/Omicron. After incubation at 37°C for 1 h, the mixtures were added to quadruplicate wells of Vero E6 cells in 96-well plates and incubated for four days. The 50% neutralizing dose (ND_50_) was defined as the highest dilution of serum that completely prevented cytopathic effects in 50% of the wells and was expressed as a log_10_ reciprocal value. HPIV3 neutralizing antibody titers were determined as described previously [41] by a 60% plaque reduction neutralization test (PRNT_60_) on Vero cells using an eGFP-expressing version of HPIV3.

### ACE2 binding inhibition assay

We investigated the breadth of the hamster antibody response by evaluating the ability of the sera to inhibit the binding of ACE2 to SARS-CoV-2 S from different strains. V-PLEX SARS-CoV-2 Panel 25 (K15586U), Panel 27 (K15609U), Panel 32 (K15671U) and Panel 34 (K15693U) kits were purchased from Meso Scale Diagnostics (MSD). The kits contain 96-well, 10-spot plates coated with S proteins from SARS-CoV-2 Wuhan-Hu-1 and variants (Alpha, Beta, Delta and Omicron sublineages). The assay was performed according to the manufacturer’s instructions and as previously described [17]. Briefly, plates were blocked and incubated with 25 μl per well of diluent or heat-inactivated sera diluted at 1:20 in diluent. Each sample was evaluated in duplicate. After 1 h incubation, 25 μl of the SULFO-TAG Human ACE2 Protein was added in each well. After additional 1 h incubation, 150 μl of MSD GOLD Read Buffer B was added in each well and plates were read using a MESO QuickPlex SQ 120MM (MSD). The electrochemiluminescence signals were analyzed by the Methodical Mind software (MSD). The ACE2 binding inhibition by each serum was shown as percent inhibition relative to the diluent.

### Analysis of host gene expression and challenge viral genome replication in respiratory tract

Total RNA was extracted from 100 μl of NT or lung homogenates using TRIzol LS Reagent and Phasemaker Tubes Complete System (Thermo Fisher) in combination with PureLink RNA Mini Kit (Thermo Fisher). SARS-CoV-2 challenge viral sgE and sgN mRNA were quantified by TaqMan-based real-time quantitative PCR, which was performed with 8 µl of RNA per reaction in triplicate wells using TaqMan RNA-to-Ct 1-Step Kit (Thermo Fisher) on QuantStudio 7 Pro Real-Time PCR System (Thermo Fisher). TaqMan primers and probes were used as previously reported [42–45] and standard curves were generated using serial dilutions of pcDNA3.1 plasmids containing each target gene sequence. The limit of detection was 3.8 log_10_ copies per g of tissue.

For quantification of host gene expression, cDNA was synthesized from 7 µl of RNA extracted as described above using the High-Capacity RNA-to-cDNA Kit (Thermo Fisher). Low-density TaqMan array cards containing primers and probes for golden Syrian hamster’s inflammatory/antiviral genes and beta-actin as the housekeeping gene are based on previous reports [46–48] and were custom-ordered from Thermo Fisher. cDNA was mixed with TaqMan Fast Advanced Master Mix (Thermo Fisher) and real-time PCR was performed in triplicate wells for each target and each sample on the QuantStudio 7 Pro. Relative expression of target genes in NTs and lungs was determined by 2^−(ΔΔCt)^ method based on the threshold cycles. Data were expressed as fold changes over the expression in NTs and lungs from six or five unimmunized unchallenged hamsters, respectively, harvested in previously published studies [17, 20]. As the expression of IFIT3 and IFN-L was not detected in any of the control hamsters, their level of expression was determined relative to the immunized/challenged hamster with the lowest detectable level of expression. The expression of IFIT3 in NT and lungs was determined relative to hamster #7 of the B/HPIV3/S-BA.5-2P group. The expression of IFN-L in NT was determined relative to hamster #3 of the B/HPIV3/S-6P+B/HPIV3/S-BA.5-2P group and in the lungs, relative to hamster #3 of the B/HPIV3 group.

### Statistical analyses

Data sets were assessed for significance using one-way ANOVA with Dunn’s post-test or two-way ANOVA with Sidak’s post-test using Prism 9.0 (GraphPad Software). Data were only considered significant at P≤ 0.05 and the exact p values are indicated in the figures. Details on the statistical comparisons are described in the figures, figure legends, and results.

## Acknowledgments

This research was supported by the Intramural Research Program of the National Institutes of Health (NIH), and by the Vaccine Research Center, an intramural division of NIAID, NIH. We would like to thank Christen Robinson, Nina Callaham, Kristina Lamot and the members of the NIAID Comparative Medicine Branch for expert support of our hamster studies. We also thank Adam S Olia, Vaccine Research Center, NIAID, NIH for providing the cDNA encoding a soluble version of the prefusion stabilized SARS-CoV-2 Omicron/BA.5 S antigen. The contributions of the NIH authors were made as part of their official duties as NIH federal employees, are in compliance with agency policy requirements, and are considered Works of the United States Government. However, the findings and conclusions presented in this paper are those of the authors and do not necessarily reflect the views of the NIH or the U.S. Department of Health and Human Services.

## Contributions

**Conceptualization:** Hong-Su Park, Yumiko Matsuoka, Ursula J Buchholz, Cyril Le Nouën.

**Formal analysis:** Hong-Su Park, Yumiko Matsuoka, Celia Santos, Eleanor F Duncan, Jaclyn A Kaiser, Lijuan Yang, Cindy Luongo, Xueqiao Liu, Ursula J Buchholz, Cyril Le Nouën.

**Funding acquisition:** Ursula J Buchholz.

**Investigation:** Hong-Su Park, Yumiko Matsuoka, Celia Santos, Eleanor F Duncan, Jaclyn A Kaiser, Lijuan Yang, Cindy Luongo, Xueqiao Liu, Cyril Le Nouën.

**Methodology:** Hong-Su Park, Yumiko Matsuoka, Celia Santos, Eleanor F Duncan, Jaclyn A Kaiser, Lijuan Yang, Cindy Luongo, Xueqiao Liu, Ursula J Buchholz, Cyril Le Nouën.

**Resources:** Hong-Su Park, Yumiko Matsuoka, Jaclyn A Kaiser, Cindy Luongo, Xueqiao Liu, I-Ting Teng, Peter D Kwong, Ursula J Buchholz, Cyril Le Nouën.

**Visualization:** Hong-Su Park, Yumiko Matsuoka, Celia Santos, Eleanor F Duncan, Jaclyn A Kaiser, Lijuan Yang, Cindy Luongo, Xueqiao Liu, Ursula J Buchholz, Cyril Le Nouën.

**Writing – original draft:** Hong-Su Park, Ursula J Buchholz, Cyril Le Nouën.

**Writing – review & editing:** Hong-Su Park, Yumiko Matsuoka, Celia Santos, Eleanor F Duncan, Jaclyn A Kaiser, Lijuan Yang, Cindy Luongo, Xueqiao Liu, I-Ting Teng, Peter D Kwong, Ursula J Buchholz, Cyril Le Nouën.

## Notes

### Competing Interest Statement

U.J.B., C.L., X.L, and C.LN. are inventors on the provisional patent application number 63/180,534, entitled Recombinant chimeric bovine/human parainfluenza virus 3 expressing SARS-CoV-2 spike protein and its use, filed by the United States of America, Department of Health and Human Services

